# A stable beneficial symbiotic relationship between endophytic fungus *Schizophyllum commune* and host plant *Panax ginseng*

**DOI:** 10.1101/175885

**Authors:** Xin Zhai, Ling Chen, Min Jia, Changhui Li, Hui Shen, Bingzhu Ye, Luping Qin, Ting Han

**Affiliations:** Department of Pharmacognosy, School of Pharmacy, Second Military Medical University, 325 Guohe Road, Shanghai 200433, China; Department of Pharmacognosy, School of Pharmacy, Shandong University of Traditional Chinese Medicine, Jinan 250355, China; School of Pharmacy, Zhejiang Chinese Medical University, Hangzhou 310053, China

**Keywords:** *Schizophyllum commune*, endophytic fungus, infection way, expression of genes, secondary metabolite biosynthetic

## Abstract

Endophytes and plants can establish specific long-term symbiosis through the accumulation of secondary metabolites. Interactions between microbial inhabitants represent a novel area of study for natural products research. In this study, a strain of endophyte 3R-2 that can enhance the biomass and contents of ginsenoside Rc, ginsenoside Rg2 and ginsenoside Rg3 of *Panax ginseng* hairy roots was screened out via HPLC, which was identified as *Schizophyllum commune* through the morphological and molecular identification. On the base, we found the infection of the endophyte were obviously observed widely in the *P. ginseng* and the strain formed a stable relationship with *P. ginseng* hairy roots in parenchyma cells around through tissues embedding slicing, HE ammonium silver staining and immunofluorescence staining. On the other hand, elicitors of fungus 3R-2 can also significantly promote hairy root growth and contents of several ginsenosides, even several times higher than 3R-2 mycelium did. Moreover, *S. commune* 3R-2 mycelium and its elicitor could enhance the transcriptional activity of key genes during the ginsenosides biosynthetic pathway dramatically. Thus, endophyte *S. commune* 3R-2 and its elicitor change the chemical substance content by regulating the expression of genes involved in the secondary metabolite biosynthetic pathway.

## Introduction

*Panax ginseng* (Araliaceae) is a traditional oriental herb that has been used to treat various diseases in East Asian countries(Mizikar 2011). *P. ginseng* has aroused great interest of researchers as a result of its different pharmacological and therapeutic effects on central nervous system, cardiovascular system and immunomodulation function(Hofseth and Wargovich 2007, Chen et al. 2008). The root of *P. ginseng* plays an important role in traditional medicine and its main component is considered to be a triterpenoid saponin of dammarane-type(Wang et al. 2001, Yue et al. 2007). As aglycones of dammarane-type triterpenes, protopanaxadiols (PPD) and protopanaxatriols (PPT) are the main part of these triterpene compounds in *P.ginseng*. As reported, Ginsenosides-Rg3, -Rd, -Rc, -Rb1, and -Rb2 were typical PPDs. The sugar moieties are connected to the ring of the triterpene dammarane at the three-position in PPD while attached at the six-position in PPT such as ginsenosides-Rg1, -Re, and -Rg2 (Sun et al. 2007). As the main PPD ginsenosides, Ginsenoside Rg2 could reduce memory impairment via anti-apoptosis (Zhang et al. 2008) and Ginsenoside Rc exhibited highest inhibitory activity against the expression of tumor necrosis factor (TNF)-α, interleukin (IL)-1β, and interferons (IFNs). (Jung et al. 2001, Choi et al. 2003). However, they are all present at low concentrations in ginseng, which is a trouble for pharmacology application (Kim et al. 2003).

Plants form various kinds of relationships with all kinds of microbes, such as mutually-beneficial symbiosis with endogenous or against pathogens in nature. Endogenous fungi are microorganisms that live within the plant’s internal tissue and do not cause any direct or obvious negative effects (Rodriguez et al. 2009). Indeed, some endophytes, such as *Piriformospora indica*, *Trichoderma atroviride* or other growth promoting endophytes can do good to plants(Ming et al. 2013). In addition, long-term colonization can lead to the accumulation of various secondary metabolites in the host(Ludwig-Müller 2015). Schulz et al. believe that the counterbalanced antagonism hypothesis can account for the relation between plants and endophytes (Schulz et al. 1999). As reported, secondary metabolites produced by host plants are likely to be the key to maintaining a balance between endophytes and host plants. Meanwhile, endogenous fungi are considered to be a powerful means to stimulate plant secondary metabolites for human use. What’s more, the effect of endogenous fungi on secondary metabolism of plants will help to target drugs through bioengineering(Ludwig-Müller 2015).

*Agrobacterium rhizogenes*-mediated transformation of hairy roots, as a quick and easy method for introducing and expressing foreign genes can be used to synthesize specific secondary metabolites in plant cells, thus providing a quick and easy method. This method not only has a higher security level, a higher growth rate, frequent branch, genetic and biochemical stability, but also improve the cosmetics, medicines and food additives content of secondary metabolites(Schulz et al. 1999). Since the growth rate of *P.ginseng* is slow, the vitro culture system was selected to obtain the indeterminate root. The hairy root culture of *P.ginseng* has become a useful platform for the research of metabolic engineering, which can replace the whole plant(Yoshikawa and Furuya 1987, Srivastava and Srivastava 2007). The present research took advantage of the platform of *P. ginseng* hairy root to study the endophyte infection ways and the function of endophytic fungus 3R-2 on *P. ginseng*, which indicated 3R-2 was a beneficial effective endophyte.

In the previous study, our research group separated the endophyte *Trichoderma atroviride* D16 from *Salvi miltiorrhiza* and found it could promote the tanshinones content obviously in access of various chemical elicitors, which gained wide attention(Ming et al. 2013). Endophytic fungus opened up a new direction for the cultivation and active substances obtaining from medical plants. As we all know, the cultivation puzzle of *Panax ginseng* (Araliaceae) and the high demand, low output of Ginsenoside Rg2, Rg3, Rc had evolved into a large restriction for pharmaceutical development. Therefore, our group isolated dozens of endopytic fungus strains and obtained an effective fungus 3R-2, which was capable of leading to an enhancement of root growth and several important ginsenosides accumulation in *P. ginseng* hairy roots. To the authors’ knowledge, there have been no reports about the effects of ginseng plant-derived endophytic fungi on the secondary metabolism of *P.ginseng*. In this study, the identification of this endophytic fungus, as well as its infection way and effects on the root growth and bio-synthesis of ginsenosides in *P. ginseng* were researched for understanding the role of endophytes in host plant survival.

## Materials and methods

### Materials and media

Fresh roots of *Panax ginseng* C. A. Mey. were collected from the city of Tonghua (JilingProvince), People’sRepublicof China. The roots were obtained from the soil and then transported to the School of Pharmacy, Second Military Medical University, Shanghai, China.

The *Panax ginseng* hairy roots used in this work were donated by School of Life Sciences in Central South University and cultured in 250 ml conical flasks including 100 ml of liquid half-strength MS medium in an orbital shaker at 25 °C and 135 rpm. The hairy roots of 1.0 g fresh weightwas culturedinevery flask for 3weeks. The furry roots were harvested in a few weeks, washed three times with distilled water, dried with paper towels, and then dried in the oven for 50°C until a constant dry weight (DW) was observed.

The reference standards of ginsenoside Rc, Rb_1_, Rb_2_, Re, Rd, Rg_1_, Rg_2_, Rg_3_ and Rh_2_ used were from Chengdu Mansite Pharmacetical CO. LTD., Chengdu (Sichuan Province), People’s Republic of China.

Potato dextrose agar (PDA) medium included 200 g potato, 20 g dglucose, 15 g agar, 1000 ml deionized water and was used to isolate and culture endophytic fungi. Whileas, 1/2MS medium contained 2.215 g MS powder, sucrose 30 g, 1000 ml deionized water and the PH was 6.0, which was applied to theculture of *Panax ginseng* hairy roots. Both types of media were sterilized for 30 min.

### Isolation and culture of endophytic fungi

The roots of *Panax ginseng* C. A. Mey. were washedthoroughly in the tap water and then the soil attaching to the root was removed with the ionized (DI) water, followed by being cut into small 15 mm- long root section with a disinfected scissors. Subsequently,those segments were surface-sterilized by using continuous immersion of 75% ethanol immersion for 30 s, sodium hypochlorite(3%available chlorine) for 3 min, 75% ethanol for 30s and finally rinsed three times in disinfectant water. After the excess water is daubed on a steroidal filter paper, the dried segments were cut into small pieces about 2mm thickness and placed evenly in a petri dish containing 100mg·l^-1^penicillin to eliminate bacterial growth.After the petri dishes were sealed with a Parafilm (Pechiney, Chicago, IL), they were cultured at 26±2 °C in the incubator until fungus growth was obivious. From the root segment, the hyphae was isolated after two or three weeks and transfered to new pure PDA. Each fungus was added to the liquid half-strength MS medium, which cultured three-week-old hairy root while the blank treatment was added with no hyphae to the fresh liquid semi-strength MS medium. In the final, the roots were collected at different intervals separately at 0, 3, 6, 9, 12 15, and 18d.

### Identification of endophytic fungus 3R-2

Endogenous fungi 3R-2grew on the PDA medium for 7 days at 28 □and photographed its morphological characteristics. The DNA of mycelium is extracted using CTAB method.According to the previous report(Ming et al. 2013), the sequence of endogenous fungi was determined according to the sequence of its 5.8 S and ITSwith primerITS 4 and ITS 5, which were compared with GenBank reference sequence. Finally, we utilized CLUSTAL X software to construct phylogenetic tree by 1000 bootstraps.

### HPLC analyses

The hairy roots were dried at 50 °C in an oven until dry weight (DW)was constant.Then these samples were ground into powder and extracted with methanol (15 mg of roots ml^‒1^) under sonication for 60 min. With the help of high-performance liquid chromatography (HPLC) system, secondary metabolites of the extract was analyzed on anAgilent-1100 system using a ZORBAX SB-C18 chromatographic column (250 mm×4.6 mm, 5 μm) at 35 °C with elution of[time(min):D(HCN)]=[0.0:19%]-[35.0:19%]-[55.0:29%]-[70.0:29%]-[100.0:40%]-[120.0:85%]. Ginsenosides of Rc, Rb1, Rb2, Re, Rd, Rg1, Rg2, Rg3 and Rh2 in the methanol extract were identifiedcompared with the available standards [ Chengdu Mansite Pharmacetical Co. Ltd., Chengdu (Sichuan Province), PR China].

### Paraffin embedding and slicing of hairy root tissues

The fresh tissues were fixed in 4% paraformaldehyde for more than 24h. After that,these tissues were removed from the fixed liquid, pruned with scissors in the fuming hood and put into the dehydration box. Then the dehydration box was placed in the basket in the gradient alcohol successively for dehydration: 75% alcohol for 4h, 85% alcohol for 2h, 90% alcohol for 2h, 95% alcohol for 1 h, anhydrous ethanol I for 30 min, anhydrous ethanol II for 30 min, alcohol benzene for 5-10 min, xylene I for 5-10 min, xylene II for 5-10 min, wax I for 1h, wax II for 1h and wax III for 1h. The immersed tissues were embeded in the melting wax on the embedding machine before it solidified.After cooling at -20 degree centigrade, wax coagulation was removed from embedding box and dressed. And the dressed wax block was cut into slices in paraffin wax slicing machine with 4 μm thickness, which werefloated and spreadon warm water at 40 degree centigrade,dried at 60 degree centigrade in oven and preserved under normal temperature.

### Ammonium silver staining and Immunofluorescence staining

The sections were put into xylene I for 20 min, xylene II for 20 min, anhydrous ethanol I for 10 min, anhydrous ethanol II for 10 min, 95% alcohol for 5 min, 90% alcohol for 5 min, 80% alcohol for 5 min, 70% alcohol for 5 min in turn and washed in distilled water. Thework preparation was blended of ammonium silver stock solution (20 ml), distilled water (15 ml) and 5% sodium tetraborate (2 ml) successively and preheated at the 60 degree centigrade in the oven. Meanwhile, these sections were placed into 1% periodic acid15 to 20 min for oxidation. Afterwards, they were rinsed with tap water for 5 min, washed with distilled water twice, circled with resistance pen, added with preheating work preparation andsealed in the wet box.After 40 minutes,the staining sections could be observed on the microscope until satisfied resultof black fungal was obtained. Subsequently, they were treated with 5% sodium thiosulfate for 2 min, rinsed intap water for 5 min, re-stained witheosin for 5 min, dehydrated through successively immersion of 95% alcohol Ifor 5 min, 95% ethanol II for 5 min, anhydrous ethanol II for 5 min, anhydrous ethanol IIfor 5 min, xylene Ifor 5 min, xylene II for 5 min, dried andsealed with neutral balsam for image collection and analysis under microscope.

The sections were putinto xylene I for 15 min,xylene II for 15 min, anhydrous ethanol I for 5 min, anhydrous ethanol II for 5 min, 85% alcohol for 5 min and 75% alcohol for 5 min and washed with distilled water.Antigen retrieval of the tissue sections were performedinboiling EDTA buffer (PH 8.0) for 5 minin the microwave.After natural cooling, the slides were washed three times (5 min each time) in PBS (PH7.4) at the decoloring shaking bed. Then the target organization was circled withresistance pen, covereduniformly with 3% BSA andsealed for 30 min at room temperature.Afterwards, the sections were dipped withprimary antibodyof certain proportion and incubated at 4 degree centigrade in wet boxovernight.Then the slides were washed three times (5 min each time) in PBS (PH7.4) at the decoloring shaking bed. After dried slightly, the sectionswere covered with secondary antibody of a correspondingspecies with primary antibody andincubated in dark at room temperature for 50 min. Then the slides were washed three times (5 min each time) in PBS (PH7.4) at the decoloring shaking bed. After dried slightly,the sections were addedwith DAPI stainingsolutionand incubated in dark for at room temperature 10 min. Then the slides were washed three times (5 min each time) in PBS (PH7.4) at the decoloring shaking bed. After dried slightly,the sections were sealed off with anti-fluorescence-quenching agent for image collection under nikon inverted fluorescence microscope.

### RNA isolation and real-time quantitative PCR analysis

3R-2 hyphae was added into the liquid half-strength MS medium with 3-week-old cultured hairy roots at a concentration of 10μl while controls were treated with fresh liquid half-strength MS medium. Hairy roots were collected at different intervals (0, 3, 6, 9, 12, 15 and 18d) and then stored at -80□. With RNA Kit II (Genebase Biosicence Co., Ltd.), total RNA was extracted from *P. ginseng*hairy root samples. The quality and concentration of RNA were examinedthrough ethidium bromide-stained agarose gel electrophoresis and spectrophotometric analysis.

Total RNA was reverse transcribed by using the RevertAid™ First Strand cDNA Synthesis Kit (Fermentas) to generate cDNA according to the manufacturer’s instruction. The gene expressions of *pgHMGR*, *pgSS*, *pgSE*, and *pgSD* were detected, respectively.The realtime PCR amplification was performed in a Lin-Gene FQD-33Adetection system (Bioer) with UltraSYBR Mixture kit (CWBIO). Each reaction included a mixture of 5 μl of SYBR Green I PCR Master Mix (ShineGene, China), 0.2 μl of forward primer (10 μM), 0.2 μl of reverse primer (10 μM),1 μl ofdiluted cDNAand 3.6 μl of RNase-free H_2_O. The reaction mixture was incubated for 10 min at 94□, and for 40 cycles of 20s at 94□, 15s at 60□ and 20s at 72□. The relative gene expression was quantified by the comparative CT method.

### Data analysis

All experimentswere analysed as their mean values and standard deviations (SD)in triplicate, including both control and different treatments of hairy root cultures, HPLC analysis, and semi-quantitative real-time PCR. The standard deviation was represented with the error barsin biological triplicates and the statistical significance of the differences was analysed by one-way analysis of variance (ANOVA) with SPASS software. Nevertheless, the statistical significance of differences in gene transcripts was analysed by one-sample t-test.

## Results

### Identification of fungal strain 3R-2

In the PDA medium, there is no sporophore and the colony is white, fluffy and the back of the colony is white too (Figure 3). Based on 3R-2 morphological characteristics and molecular analysis of ITS rDNA (ITS 1, ITS2, and 5.8 S rRNA genes), the fungus had been identified as *Schizophyllum commune*. The acquired ITS-5.8 S rDNA sequence has been deposited as assession number KU042974 in GeneBank. With the neighbor-joiningmethod, we carry on phylogenetic tree after 1000 bootsrap and identification at the species classification level is based on ≥ 100% similarity (Figure 4).In addition, the plant endogenous fungi have been collected and deposited in the Chinese General Culture Collection Center(CGMCC) in Beijing, China as 11009 (Figure 3).As we know, there is no report that *Schizophyllum commune* was able to increase plant growth and active substances.

**Figure 3.**
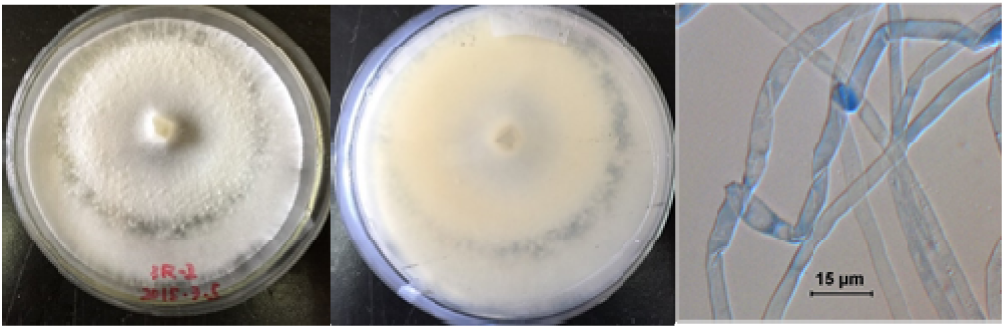
The morphology of endophytic fungi 3 R-2 mycelium on PDA (A) Colony positive photo, (B) Colony opposite photo and (C) Micrograph of 3R-2.

**Figure 4.**
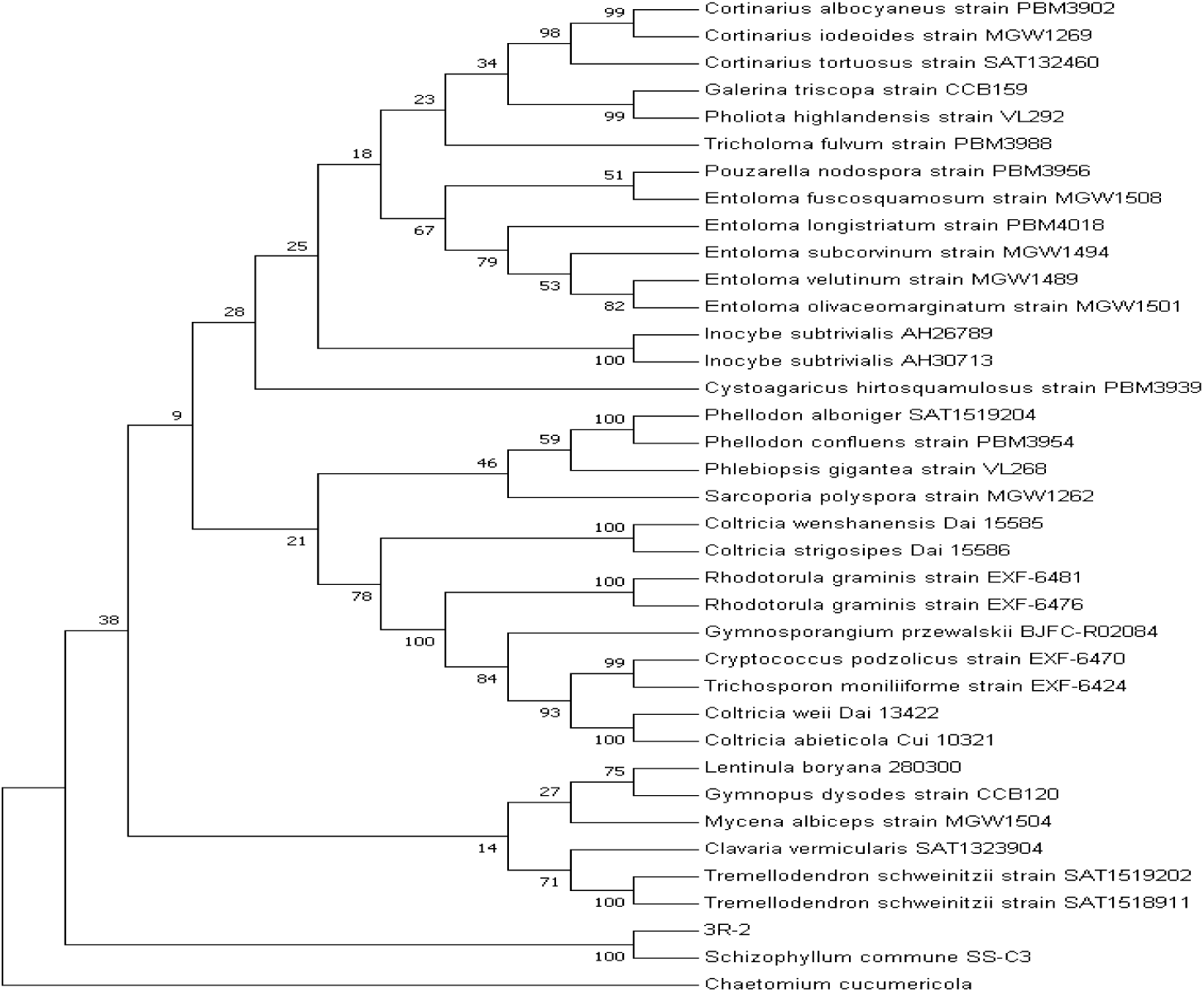
The phylogenetic tree of the endophytic fungus 3R-2, *Chaetomium cucumericola* is used as outgrout

### The observation of 3R-2 mycelia in P. ginseng hairy roots

The existence of endophytes and the morphology of endophytes were obviously observed in *P. ginseng* hairy roots through *P. ginseng* tissues embedding slicing, HE ammonium silver staining and immunofluorescence staining. Compared with 4R-2 strain, more 3R-2 mycelia inoculated in *P. ginseng*hariy root and 4R-4 had the same inoculation capacity with 3R-2, whereas 4R-4 constructed hyphae web throughout root cells(Fig. 6). HE ammonium silver staining illustrated the main infection sites were found in parenchyma cells around (Fig. 7). HE ammonium silver staining represented the infection stable state. Significantly, intracellular hyphae remained staying in parenchyma cells and enveloped by host cell membrane,which was for longer periods not just for infection. The means of endophytic fungi infection on plant tissues may provide some experimental basis for the future research.

**Figure 6.**
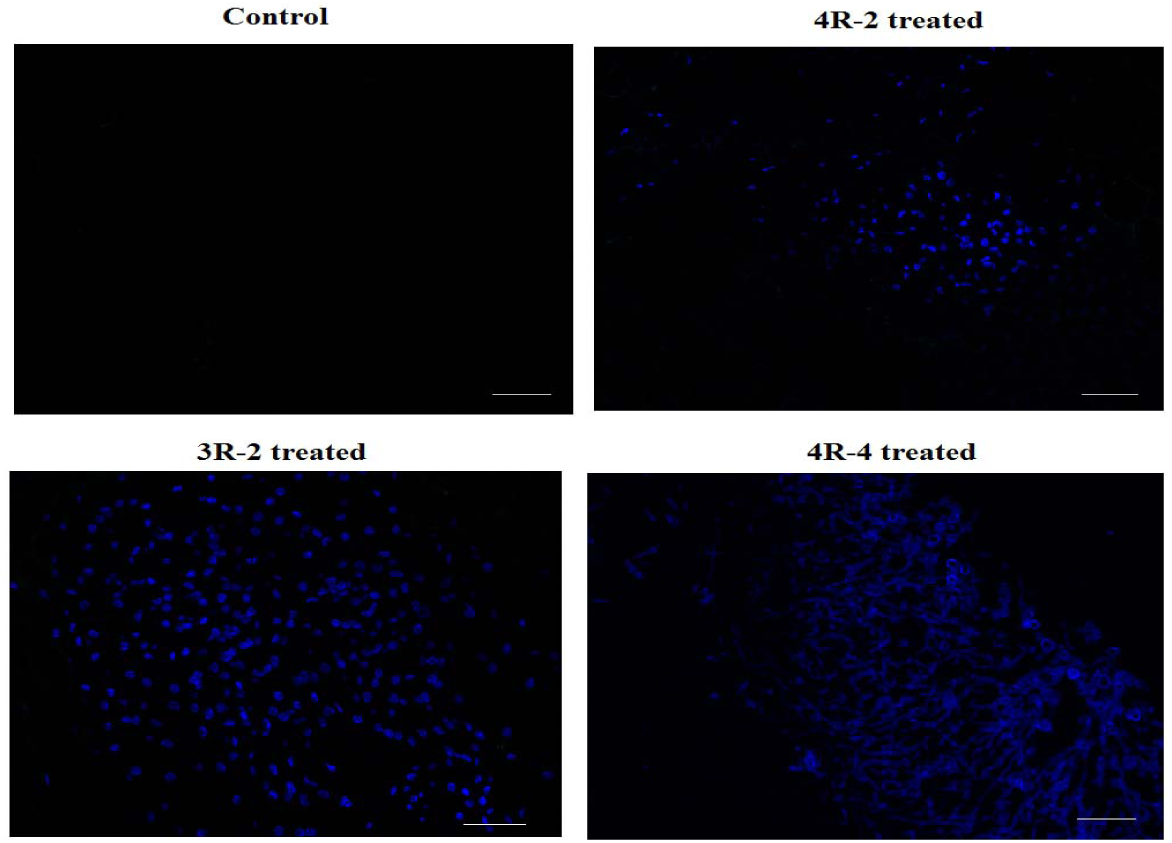
Different endophytic fungi in *P. ginseng* hairy roots after immunofluorescence staining (blue-fluorescence represented the mycelia, amplification factor:400×)

**Figure 7.**
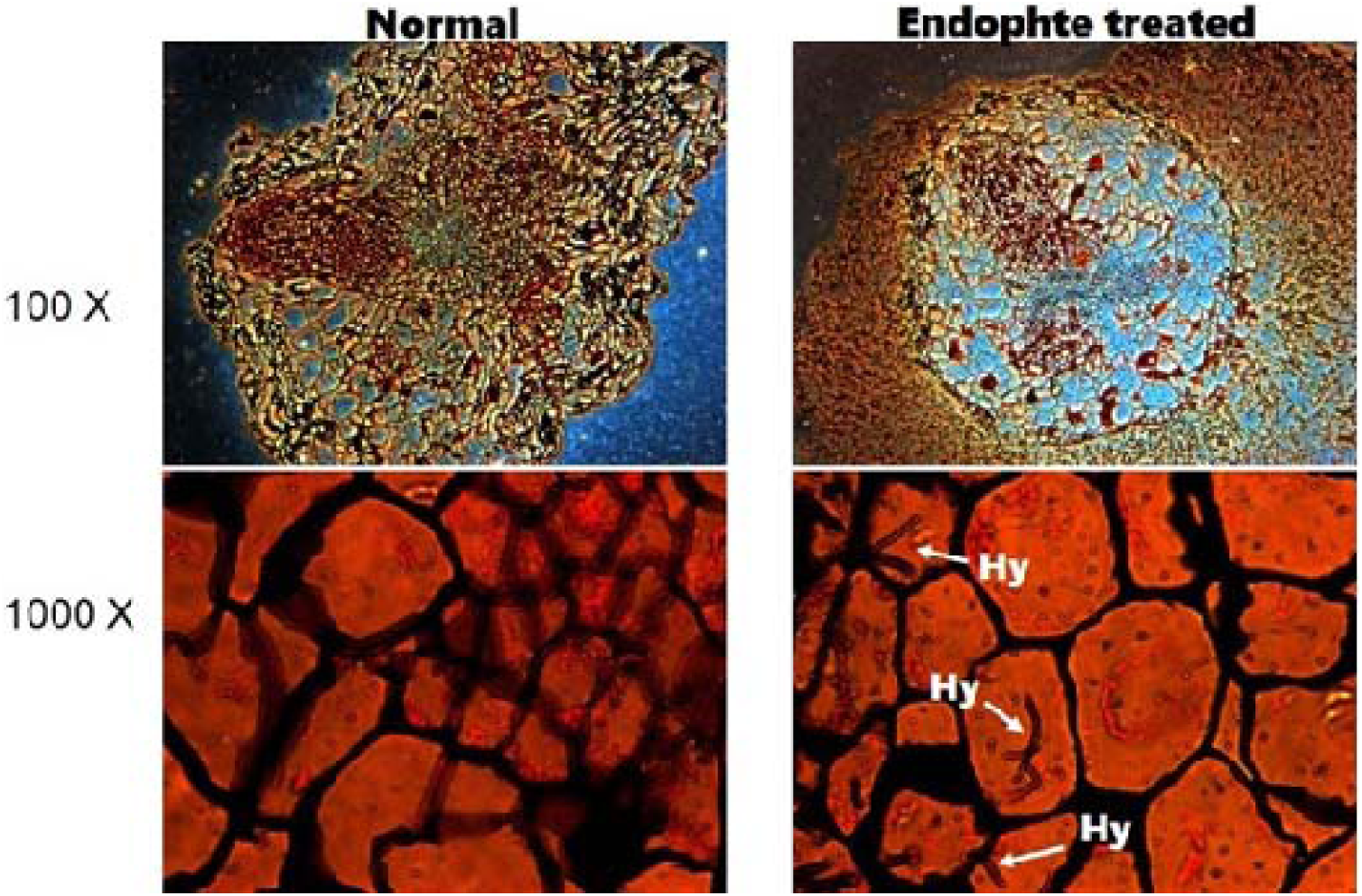
The endophytic fungi *P. ginseng* hairy roots after ammonium silver staining (the arrows indicate the mycelia)

**Figure 8.**
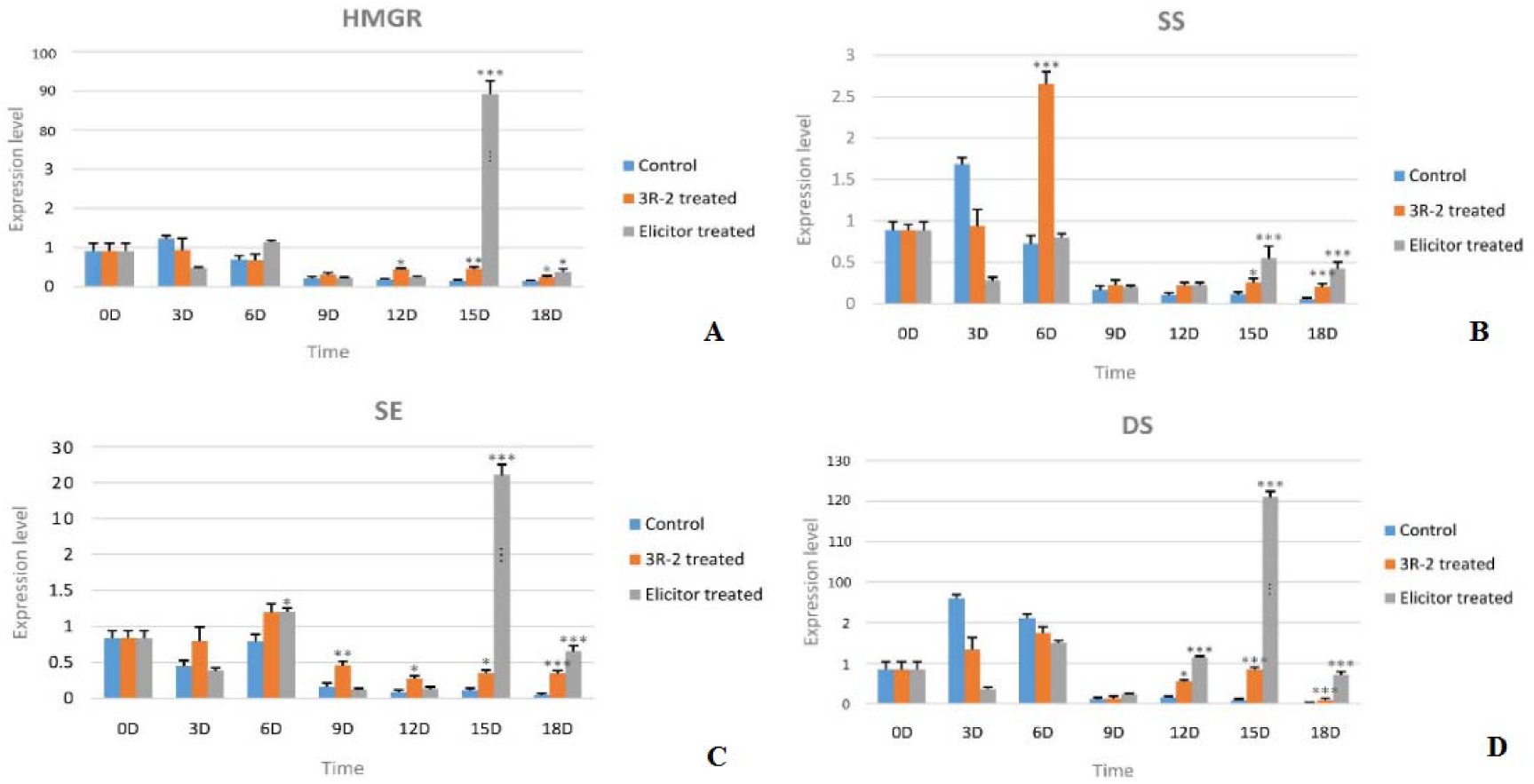
Effects of endophyte 3R-2 and elicitor from 3R-2 on the expression of genes in the ginsenoside biosynthetic pathway in *p. ginseng* hairy roots. (A) HMGR, hydroxymethylglutaryl-CoA reductase; (B) SS,squalene synthetase; (C)SE, squalene epoxide enzyme; (D) DS,darnmarenediol synthase.(*:*P*<0.05, **:*P*<0.01, ***:*P*<0.001 vs control;n=3)

### Effects of 3R-2 mycelium on the biomass and ginsenoside contents of P. ginseng hairy roots

Eighty three endophytic fungal strains were isolated from the roots of *Panax ginseng*on the basis of morphology. A strain of endophyte 3R-2 that could increase the contents of ginsenoside Rc and ginsenoside Rg_2_ was screened out by using HPLC. Compared with other endophytic fungi, 3R-2 has exceptional effect on the growth and bioactive substances accumulation of *P. ginseng* hairy roots(Fig. 1). Endophyte 3R-2 can promote biomass of *P. ginseng*hairy roots by 16% and the content of Rg_2_ and Rc were 11.74 fold and 2.75 fold seperately under the treatment of 3R-2 in comparison with control group, which indicated 3R-2 was an effective beneficial endophyte for *P. ginseng*.

**Figure 1.**
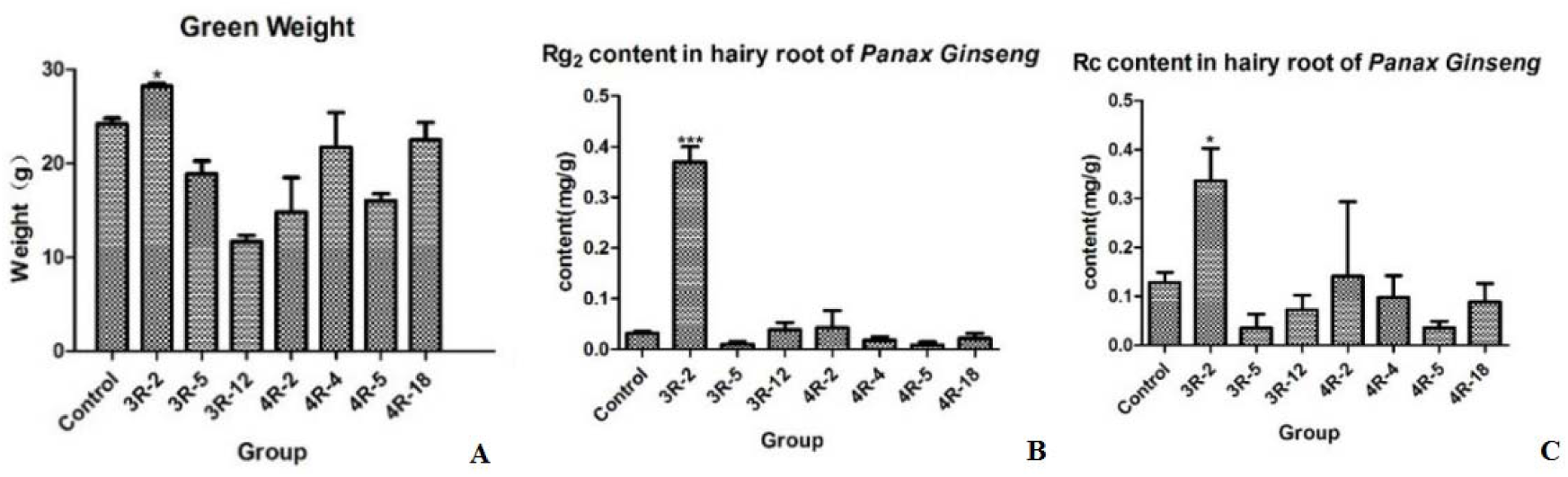
Effects of 3R-2 mycelium on the biomass, Rg_2_ content and Rc content of *P. ginseng* hairy roots (*: *P*<0.05, **: *P*<0.01, ***: *P*<0.001 vs control;n=3)

The endophyte 3R-2 hyphae was added into of *P. ginseng* hairy root cultures by means of punching, the effects of endophyte 3R-2 hyphae on the growth and secondary metabolic biosynthesis of *P. ginseng* hairy roots were studied.Results show that the endophyte 3R-2 has no obvious effects on the biomass and contents of ginsenoside Rc and ginsenoside Rg_2_ of *P. ginseng* hairy roots within seven days, whereas it significantly promoted the biomass and contents of ginsenoside Rc, ginsenoside Rg_2_ and ginsenoside Rg_3_ of *P. ginseng* hairy roots within 14-21 days; the content of ginsenoside Rc reached the highest in 14^th^ days after endophyte 3R-2 hyphae co-culture with *P. ginseng* hairy roots (Fig. 2).

**Figure 2.**
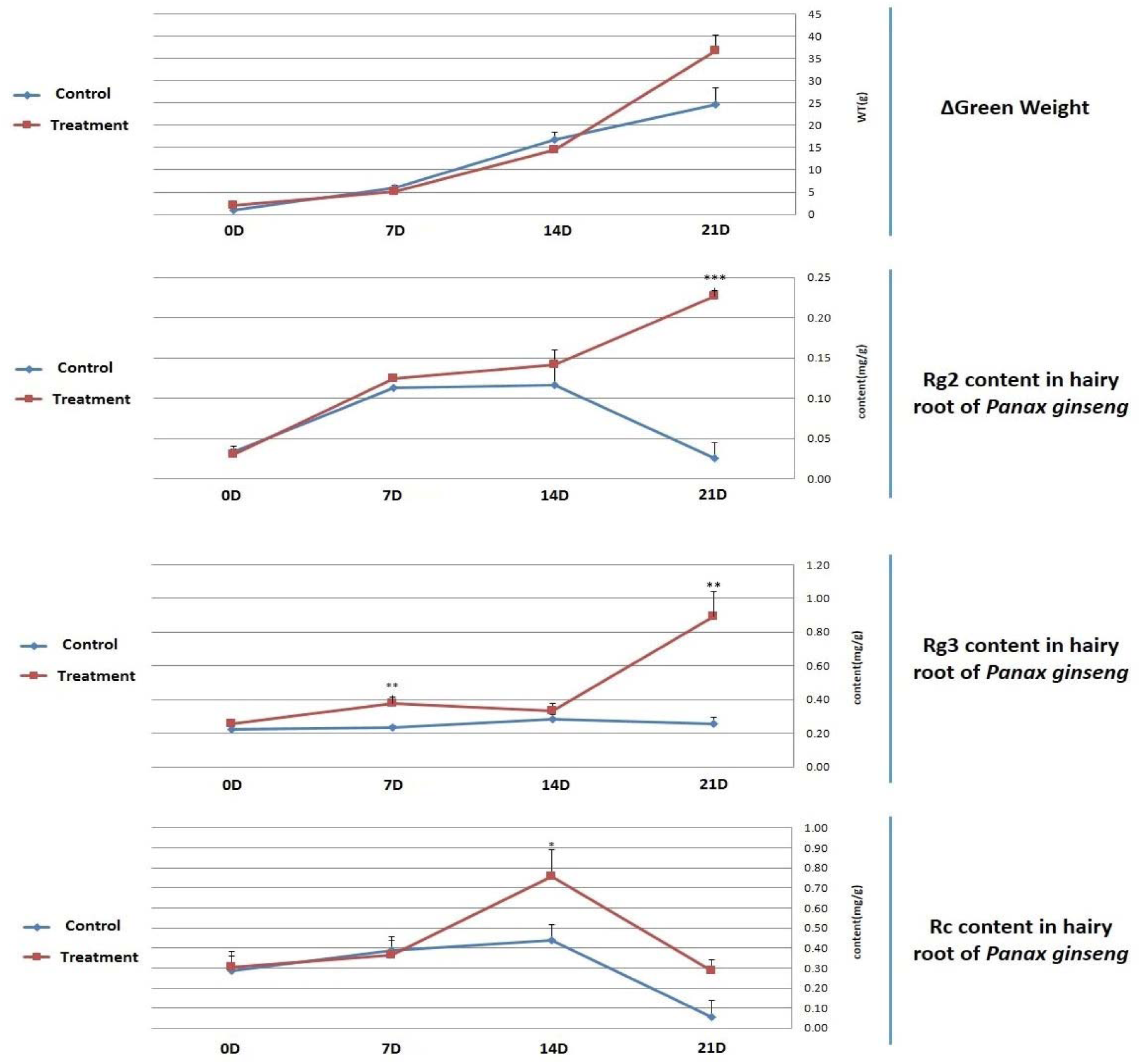
Effects of 3R-2 mycelium on the green weight and biosynthesis of Rg_2_, Rg_3_ and Rc in *P. ginseng* hairy roots (*: *P*<0.05, **: *P*<0.01, ***: *P*<0.001 vs control;n=3)

### Effects of Schizophyllum commune on the expression of genes in the ginsenoside biosynthetic pathway in P. ginseng hairy roots

The RT-PCR of four key enzyme genes HMGR, SS, SE and DS in the secondary metabolic pathways of ginsenosides was carried out, and the results showed that key enzyme genes SS of the 3R-2 mycelium group is significantly higher than the control group and reached the highest in 6 days, which promoted the expression of SS by 3.86 fold in 6 days. From 12^th^ to 18^th^ day, 3R-2 mycelium had an obvious effect on increasing the expression of HMGR and DS (Fig. 7A,D). And the expression of SE was increased from 9^th^ to 18^th^ day (Fig. 7C), whereas 3R-2 mycelium cause a distinct boost on the expression of SS from 15^th^ to 18^th^ day(Fig. 7B). While the key enzyme genes HMGR, SE and DS of the 3R-2 elicitor groups reached its highest in 15 days and were 90 times, 20 times and 120 times higher than control group, respectively (Fig. 7). The elicitor of 3R-2 promoted the expression of HMGR in the late days from 15^th^ to 18^th^ day. Nevertheless, 3R-2 elicitor had a tremendously greater increase on the expression of HMGR, SS, SE, DS by tens times.

## Disscusion

Ginseng has been known for thousands of years in the Far East as a precious medicinal herb. In recent years it has attracted interest in western countries. It was widely used and extracted for pharmacological and therapeutic usage, and now restricted by low output of *P. ginseng* root and low content of ginsenosides. In this study, endophytic fungi from *P. ginseng* were chosen as research objects to study the relationship with the host plant, aiming to promote ginseng growth.. We adopted the *P. ginseng* hairy root system as the research platform and screened out an endophyte 3R-2, which could promote the growth of host plants and accumulate the effective ginsenosides contents in *P. ginseng* hairy roots. Through the morphological and molecular identification, the target endophyte was identified as *Schizophyllum commune.* There are some reports about endophyte *Schizophyllum commune*, which is a common “miscellaneous fungus” all around the world, especially in tropical and subtropical miscellaneous tree forests. *Schizophyllum commune* was an vigorous strain and mycelium extracts contain active substances, exhibiting obvious inhibitory effects on *Staphylococcus aureus*, *Escherichia coli*, *Dysentery bacilli*, *Bacillus subtilis* and *Salmonella paratyphi B* as reported(Wang et al. 2001). Endophyte *Schizophyllum commune* not only promoted the growth of *P. ginseng* hairy root but also increased contents of ginsenoside Rc and ginsenoside Rg2, which indicated *Schizophyllum commune* 3R-2 was an efficient endophytic fungus for *P.ginseng*, and the active substance of 3R-2 could be a great study point (Kei et al. 2016, Jaber and Enkerli 2017).

Through tissues embedding slicing, HE ammonium silver staining and immunofluorescence staining, we found the endophytes were obviously observed in fresh roots, stems, leaves and fruits of cultivated *P. ginseng*. The mycelium in leaves and fruits of *P. ginseng* observed under the microscope is the most obvious, while endopyhtic fungi were distributed in root and stem sparsely (Fig.5). As researched, the hyphae should break through the root periderm from soil at the first time and then enter into the epidermal cells, which represented the interaction between mycelia and live plant cell(Kei et al. 2016). Subsequently, endophyte may transfer and spread through vascular bundles to stems, leaves and fruits (Sesma and Osbourn 2004). As revealed, the environment of fruits and leaves was more favorable to endophtic fungi survival. After that, the infection situations of several strains important endophytic fungi in *P. ginseng* hairy root tissues were observed by using the same method (Fig.6). Compared with 4R-2 strain, more 3R-2 mycelia inoculated in *P. ginseng* hariy root, which indicated that 3R-2 was more inclined to colonize the root and develop a biotrophic relationship with the *P.ginseng* (Fig. 6). Obviously, 4R-4 had the same inoculation capacity with 3R-2, whereas 4R-4 constructed hyphae web throughout root cells(Fig. 6). The 4R-4 manisfested obvious aggressiveness than 3R-2 and did not enhance the content of ginsenoside Rc, Rg2 and Rg3 (Fig. 1), which illustrated reciprocal stable colonization state estabolished on less offense and more reside. To study the inoculation means, HE ammonium silver staining was applied into 3R-2 colonization and it signified that parenchyma cells was the main resident interactional place (Fig. 7). Intracellular hyphae lived in parenchyma cells and was enveloped by host cell membrane for longer periods (Bonfante and Genre 2010). As author’s knowledge, it was the first time to investigate the infection way in *P. ginseng*.

**Figure 5.**
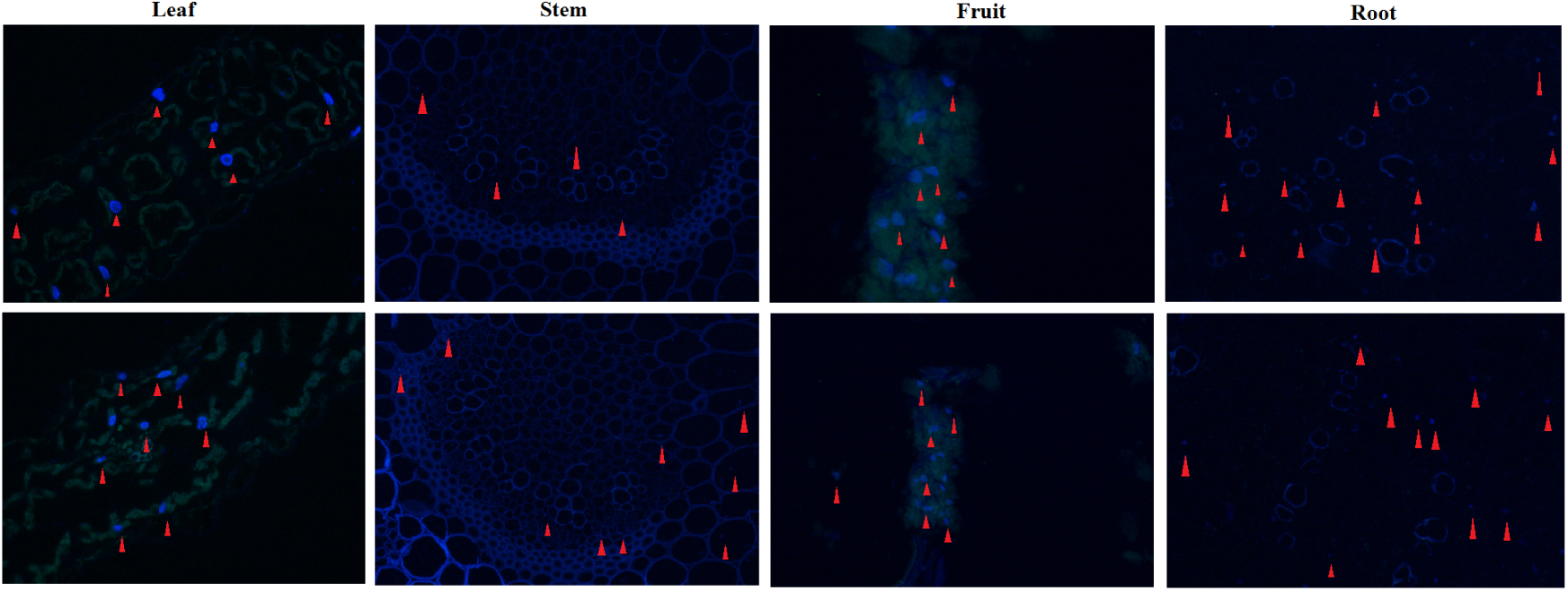
Endophytic fungi distribution in different tissues of *P. ginseng* after immunofluorescence staining (blue-fluorescence and red triangle represented the mycelia, amplification factor:400×)

In our following study, the biomass of *P. ginseng* hairy roots were tested and it was promoted by endophyte 3R-2 greatly. While most studies on the interactions between plants and endophytic fungal have so far focused on the benefits of such interactions to host plants through increased tolerance and resistance to diseases, only a handful of studies to date have investigated the potential role of endophytic fungal as plant growth promoters(Elsharkawy et al. 2012). Exploring the full potential of interactions between plants and fungal could facilitate a more effective use of these fungi for biocontrol strategies, which could present possible explanations for the lack of consistency in the plant growth promotion obtained by the inoculation of fungi(Chandanie et al. 2009). On the 21^th^ day after inoculation of 3R-2, the biomass of *P. ginseng*hairy roots was increased by1.6 fold, which has revealed its potential for application as a plant growth-promoting mycorrhizal fungus for realizing the targeted improvement in the production of medical plants *P. ginseng*. According to the reports, endopyhtic fungi ameliorated the plant growth by secreting gibberellins(GA), indole 3-acetic acid (IAA) (Abdul et al. 2012) and other signalling molecules generall (Straub et al. 2013)(Cavalcante et al. 2007) or increased the photosynthesis efficiency (Harman 2011). It was considerably difficult for endophytic fungi to form plant-fungus symbiosis(Rodriguez and Redman 2008), let alone colonize in the root for a long period to increase the growth of plant(Bae et al. 2009). Only highly effective strains can open the entrance door to initiate colonization with the invertase key (Vargas et al. 2009) and interact with the host through sucrose-independent network and chemical communicants releasing (Shoresh et al. 2010, Vargas et al. 2011). *P. ginseng* was hard to build symbiotic association with fungi, which illustrated *Schizophyllum commune* 3R-2 was a highly efficient strain for *P. ginseng.* As to the growth-promoting mechanism, it can be researched further in the next experiment.

And the change of the active ingredients in *P. ginseng* hairy roots were tested by high performance liquid chromatography technique during the whole coculturing period. The contents of ginsenoside Rc, ginsenoside Rg_2_ and ginsenoside Rg_3_ of *P. ginseng* hairy roots were greatly enhanced within 14-21 days and the content of ginsenoside Rc reached the highest in 14^th^ days after endophyte 3R-2 hyphae co-culture with *P. ginseng* hairy roots. When 3R-2 was inocubed into *P.ginseng* hairy root, the hyphae may have to adapt to the cultivated condition through several days. After adaptation phase, 3R-2 gradually interacted with *P.ginseng* hairy root and form a beneficial mutualistic relationship. The ginsenosides may be one of interaction factors and metabolic flow was strengthen in order to reach a balanced system(Berg et al. 2016). Finally, the change of key enzyme genes in ginsenosides metabolic pathways were also tested by RT-RCR techniques. We expect to provide some experimental basis and scientific evidence to reveal how endophytic fungi affect quality of Chinese medicinal plants *P. ginseng* in the preliminary study(Ghaffari et al. 2016). Ginsenosides belong to triterpenoid compounds and of the biosynthetic pathways of them have been preliminary studied, which can be divided into three stages: (1) The biosynthesis of IPP and DMAPP; (2) Biosynthesis of 2, 3-oxidosqualene; (3) oxidation of oxidosqualene and modification of the complex functional groups. In order to further analysis the influence of 3R-2 mycelium on the ginsenosides biosynthesis, we selected the four key enzymes genes HMGR, SS, SE and DS of ginsenosides biosynthetic pathways in RT-PCR studies. These key enzyme genes expressions of ginseng hairy roots were quantitatively analyzed when cocultured with endophyte 3R-2 in 0, 3, 6, 9, 12, 15 and 18 days. In this study, the results were shown in fig. 6. Strain of 3R-2 significantly promoted the expression of gene SS in the 6th days, and significantly promoted the expressions of HMGR, SE and DS in the 15th days. The strain of 3R-2 promoted the expressions of these key enzymes, which were consistent with the results of HPLC. On the other hand, in order to remove the virulence of 3R-2, we investigated the activity of the fungi elicitors, and the effect was several times higher than 3R-2 mycelium.

Similarly, we also obtained the same results in metabonomics studies. A large number of compounds were found greatly increased except the several increased ginsenosides in HPLC test(Sun et al. 2009). Next step is to identify these compounds and establish the compound libraries for the purpose of providing the basis for the production and application in the future. As studied, the results suggested that *Schizophyllum commune* could cause chemical defence responses in the host when confronting pathogens(Tenkanen and Siikaaho 2000, Tan and Zou 2001). This endophyte changed the content of chemical composition in the host plant by affecting the expression of genes related to the secondary metabolite biosynthesis pathway. As a result, the addition of *Schizophyllum commune* can be considered as an effective approach for the large-scale production of ginsenosides in *P. ginseng* hairy root culture systems or exploit 3R-2 function to manufacture into biotic fertilizer for high production and superior quality (Tkacz and Poole 2015). However, the active substances responsible for the stable relationship between *Schizophyllum commune* and *P. ginseng* still keeps unclear and the reason of promoting hairy root growth and stimulating the biosynthesis of ginsennosides in the hairy root culture by *Schizophyllum commune*t will be clarified in future studies.

